# Surf2Spot: A Surface-Informed Geometry-Aware Model for Predicting Binder and Nanobody Design Hotspots

**DOI:** 10.64898/2026.01.23.700806

**Authors:** Anwen Zhao, Yu Li, Tianhao Wu, Ran Fu, Zixuan Wang, Yanfen Xu, Xuliang Han, Yuxuan Lou, Wenying Wang, Jun Yan, Xiangfeng Wang

## Abstract

Protein-protein interactions (PPIs) and nanobody-antigen interactions (NAIs) play essential roles in cellular function, yet accurate hotspot prediction remains challenging. We present Surf2Spot, a deep learning framework that integrates sequence embeddings, structural features, and protein surface properties to predict interaction hotspots. By jointly modeling structural and physicochemical determinants, Surf2Spot achieves superior performance on curated PPI and NAI datasets, improving F1-scores by 45.9-46.9% and AUPRC by 43.2% over existing methods. Case studies on NbPDS1 and VdPDA1 demonstrate that Surf2Spot accurately recovers experimentally validated hotspots clustered within functional domains. When incorporated into binder design pipelines (RFdiffusion and BindCraft), Surf2Spot-guided designs yielded a 4-fold increase in successful design throughput and enhanced binding affinities compared to baseline strategies. These results establish Surf2Spot as a powerful tool for hotspot discovery and rational protein engineering.

## Introduction

Protein-protein interaction (PPI) is a fundamental process underlying cellular activities such as signal transduction, transport, and metabolism^1^. Accurate identification of the structural surfaces and residue-level hotspots that mediate PPIs is critical for understanding disease and for engineering molecular tools in therapeutic development and biotechnology. These interaction hotspots often contribute disproportionately to the total binding free energy of a protein–protein interaction, despite comprising only a small subset of surface residues, and their accurate identification is essential for structure-based approaches to modulating protein interfaces.

Recent advances in generative modeling have revolutionized protein design, particularly through the development of diffusion-based frameworks that generate binding scaffolds followed by inverse folding^2^. Notable examples include RFdiffusion for plausible backbone generation coupled with ProteinMPNN for *de novo* sequence design^3,4^, BindCraft for integrated sequence-structure design workflows^5^, and IgGM for antibody and nanobody modeling^6^. All of these methods rely on an initial definition of “hotspot” residues within the target surface to initiate the diffusion process^7,8,9^. These hotspots act as anchors for guiding binding site generation, and their accurate identification directly impacts the success of downstream binder design.

In ideal cases, where a reliable co-crystal structure of the target protein complex is available, hotspots can be extracted from known interfaces^10,11^. However, this approach is severely limited by the availability of experimental structures^12^, as only about 50% of structures in the Protein Data Bank represent protein-protein complexes^13^. As a result, there remains a pressing need for methods that can predict binding hotspots from unbound or predicted monomeric structures.

Several computational tools address this challenge. Structure-based methods like HotPoint^14^, KFC2^15^, PredHS^16^, and PredHS2^17^ use energy-based scoring or mutational scanning but depend on known complexes. Sequence-based methods such as SPOTONE^18^, and structure-aware tools like PPI-hotspot^ID19^ do not require a complex structure but still rely on evolutionary features or native folds. Epitope prediction tools used in antibody design, such as SEMA^20^ and DiscoTope^21^, also attempt to identify contact-prone surface residues, but they are limited to antibody–antigen contexts. These methods, while useful, often lack generalizability to novel, uncharacterized proteins or targets with minimal structural or evolutionary data.

With the emergence of proteolysis-targeting chimera (PROTAC) strategies^22,23, 24,25^, and their biological equivalents(bioPROTACs^26,27^), there is increasing interest in designing modular binders capable of targeting endogenous proteins for degradation. Nanobodies, in particular, offer several advantages due to their small size, stability, and capacity to recognize flat or cryptic epitopes^28,29^. However, hotspot prediction remains a bottleneck for applying such strategies in non-model organisms, especially given the scarcity of specific structural and interaction data.

To address this need, we developed Surf2Spot, a computational tool that predicts interaction hotspots directly from a protein sequence or structure. Built on a dynamic graph convolutional neural network (DGCNN) model^30^, Surf2Spot is trained on both protein–protein and nanobody–antigen interaction data to generate two complementary hotspot classes applicable to binder or nanobody design^31^. In contrast to previous methods^18,19^, Surf2Spot uses a point cloud-based representation of protein surfaces, capturing both geometric and physicochemical features while explicitly excluding buried residues unlikely to participate in binding^30^. This architecture enables Surf2Spot to operate effectively on predicted monomeric structures and enhances hotspot interpretability and downstream design success—particularly in targets where structural data are sparse.

## Results

### Surf2Spot: A Surface-Aware Graph Neural Network for Hotspot Prediction of Monomeric Proteins

We developed Surf2Spot, a deep learning framework that predicts protein– protein interaction (PPI) and nanobody–antigen interaction (NAI) hotspots using only monomeric sequence and structure. Unlike prior methods that rely on co-crystal complexes, Surf2Spot leverages a surface-aware graph representation built from predicted 3D structures and point cloud features **(Fig. 1a)**. Each input protein sequence is first processed by two pretrained models: ProtTrans^32^, which generates informative sequence embeddings, and ESMFold^33^, which predicts the 3D structure. From the predicted structure, we extract residue-level properties, including secondary structure, torsion angles, and relative solvent accessibility, using DSSP^34^. These are combined with one-hot amino acid encoding and a spatial distance featurizer that generates adjacency matrices based on residue proximity^35^ **(Fig. 1b)**. To manage challenges associated with long sequences, such as data imbalance and high spatial complexity^36^, we employed two distinct but complementary approaches: Chainsaw segments proteins exceeding 400 AAs into subdomains to manage structural complexity^37^, while GPSite scans complete structures for high-probability interaction residues^38^ **(Fig. 1a)**.

**Fig. 1:**
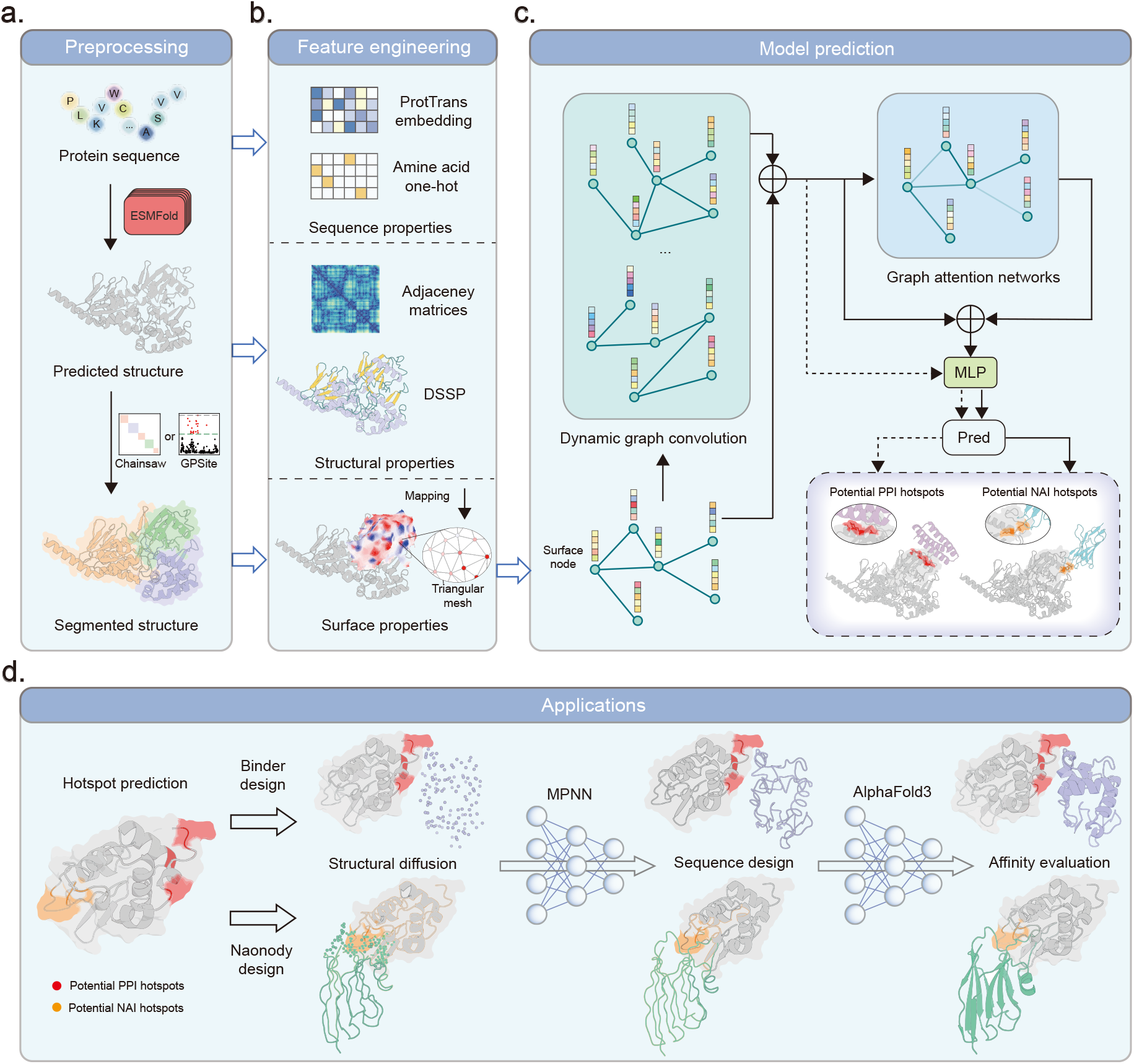
Overview of the Surf2Spot pipeline. **a**, The input protein sequence is processed by the pre-trained language model ProtTrans and the folding model ESMFold to generate sequence embeddings and predicted structures, respectively. For oversized structures, we use domain segmentation or PPI probability enrichment for segmentation. **b**, One-hot encoding is constructed from this sequence, while DSSP-derived secondary structure properties and the residue adjacency matrix are extracted from the predicted structure. We apply the MaSIF-based method to sample and extract surface features (shape index, Poisson-Boltzmann continuum electrostatics, hydrogen-bond potential, and hydropathy) and to generate a triangular mesh of surface point clouds. ProtTrans embeddings and adjacency matrix features are projected into lower-dimensional spaces. These features are aligned via Euclidean distance-based mapping and integrated with the surface point cloud to form a graph structure. **c**, This graph is then input into a three-layer DGCNN and, for nanobody-antigen interaction (NAI) hotspots, a GAT network. Protein-protein interaction (PPI) hotspots omit the GAT step (dotted path). Residual connections are applied throughout. A final MLP predicts hotspot likelihood. Applications of Surf2Spot include hotspot identification for both PPI and NAI. **d**, Surf2Spot facilitates key downstream applications in protein design by accelerating the design-build-test cycle, thereby streamlining workflows across biology, biotechnology, and synthetic design.

The protein surface is represented as a graph using the MaSIF pipeline^39^: surface points are sampled and encoded with physicochemical properties (e.g., shape index, hydrogen-bond potential, hydropathy, Poisson-Boltzmann continuum electrostatics)^40^, then connected by a triangular mesh^41^. Residue-level features are mapped to these surface nodes via Euclidean distance, producing a surface graph with rich node and edge attributes, including residue distances, sequence embeddings extracted by ProtTrans, structural features constructed by DSSP, and surface features sampled with MaSIF **(Fig. 1b)**. These graphs are processed using a dynamic graph convolutional neural network (DGCNN)^30^. For NAI prediction, an additional graph attention network (GAT) captures long-range spatial context **(Fig. 1c)**.

Surf2Spot processes heterogeneous features via dedicated MLP encoders to generate initial graph embeddings^42^, which are passed through a three-layer DGCNN module with edge recomputation at each step. For NAI prediction, an additional three-layer GAT module operates directly on the original point cloud topology to capture long-range dependencies^43^. Residual connections are applied throughout, culminating in a fully connected layer that outputs residue-wise hotspot probabilities. This architecture culminates in a fully-connected layer that transforms the final graph representation into hotspot probability predictions **(Fig. 1c)**. These predicted hotspots can directly inform the design of binders and nanobodies, providing actionable insights for targeting specific epitopes **(Fig. 1d)**.

Surf2Spot distinguishes itself from conventional approaches through three fundamental innovations: (1) it bypasses multiple sequence alignments (MSAs) and native structures^44^ by using ProtTrans and ESMFold, enabling large-scale, low-cost prediction^44^; (2) it predicts interaction sites from unbound structures, avoiding limitations imposed by simplex binding partners; and (3) its surface-based graph construction inherently excludes buried amino acids, improving hotspot balancing and structural robustness. Together, these features make Surf2Spot particularly well suited for practical applications in binder and nanobody engineering, where rapid and accurate epitope identification is essential.

### Task-Specific Dataset Design and Model Validation for Hotspot Prediction

To train Surf2Spot for both PPI and NAI hotspot prediction, we curated two distinct datasets reflecting their biological differences. PPI hotspots, traditionally defined as residues whose alanine substitution reduces binding energy by >2 kcal/mol^45,46,47,48,49,50,51^, were sourced from 507 non-redundant complexes (filtered from 1,334 complexes from PDBbind2020 and GPSite^52^). NAI hotspots, by contrast, reflect epitope functionality and are enriched in charged, hydrophilic residues (e.g., Arg, Asp, Glu), requiring distinct modeling from the hydrophobic-dominated PPI hotspots (e.g., Leu, Tyr, Phe) **(Fig. 2a)**^53^. A total of 248 high-quality nanobody– antigen complexes were selected from 1,248 entries from the SabDab database, using identical structural filters^54^. Notably, both PPI and NAI hotspots were more frequently distributed on protein surfaces compared to linear sequences, reflecting the spatial clustering of functional residues within tertiary structures **(Fig. 2b-c)**^53^. Furthermore, residue conformations differed markedly between monomeric and complexed states, with the most pronounced discrepancies occurring at the interface **(Fig. 2d-e)**. To avoid overfitting to complex-bound conformations, all samples were supplemented with ESMFold-predicted monomer structures, and each complex-monomer pair was co-assigned to training or test sets for unbiased evaluation. This surface-centric approach focuses on binding interface features, enhancing prediction accuracy and efficiency.

**Fig. 2:**
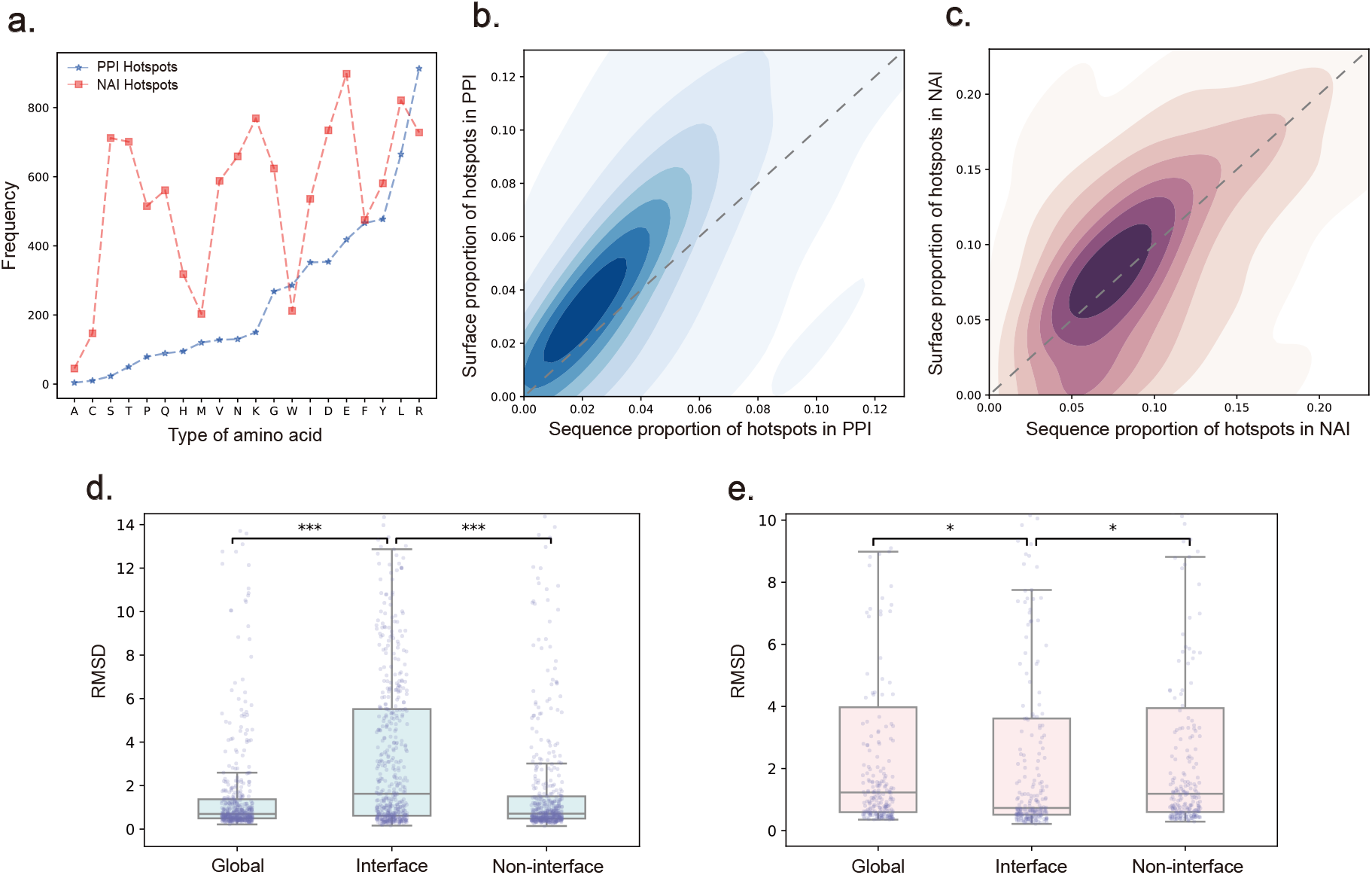
Dataset construction and training of Surf2Spot. **a**, Amino acid frequency distributions in PPI and NAI hotspots. Frequencies of the 20 amino acids are plotted in ascending order of PPI frequency. **b, c**, Distribution of hotspot residues by surface and sequence in the PPI and NAI datasets, respectively. Contour plots show the density of surface-to-sequence hotspot ratios normalized per protein. The dashed line (y = x) represents equal distribution. **d, e**, Comparison of residue RMSD between ESMFold-predicted and experimental structures for the (d) PPI and (e) NAI datasets. The binding interface is defined by a Euclidean distance cutoff of 5 Å for PPI and 8 Å for NAI from the partner nanobody. Significance codes: ***p < 0.001, **p < 0.01, *p < 0.05. **f, g**, Final model performance on the PPI and NAI datasets shown as radar plots of AUCPR, precision, recall, F1-score, and MCC.

We applied 5-fold cross-validation to assess model performance for predicting PPI and NAI hotspots. Each fold trained a DGCNN model with the Adam optimizer (learning rate = 1e-4, weight decay = 1e-5) and a learning rate scheduler. Metrics including accuracy, precision, recall, F1, Matthews correlation coefficient (MCC), and ROC-AUC were monitored throughout training, with F1-based thresholds selected for final evaluation. Results from all folds were aggregated to evaluate generalization. Surf2Spot achieved high predictive performance on both tasks: F1 scores of 0.610 and 0.507, and MCC values of 0.602 and 0.453, respectively **(Supplementary Fig. 2)**. These results demonstrate robust generalizability across diverse binding modalities and highlight the value of surface point cloud modeling, which excludes buried residues and enhances hotspot signal quality.

### Surf2Spot Outperforms Leading Hotspot Prediction Methods

We benchmarked Surf2Spot against two state-of-the-art tools for hotspot prediction on unbound protein structures: the sequence-based SPOTONE and the structure-based PPI-hotspot^ID^. All models were evaluated on the same test set (Test_52) using both global amino acid-level and per-protein average evaluations. Hotspot prediction is inherently imbalanced, making metrics like F1-score, MCC, and AUPRC more informative than accuracy or AUROC^55^. Surf2Spot vastly outperformed both comparators across these metrics, indicating better balance between precision and recall and more robust binary classification performance, yielding a 45.9% relative improvement in F1-score and more than doubling the MCC in global evaluation (**Table S1-1, S1-2**). While PPI-hotspot^ID^ and Surf2Spot showed comparable performance in AUROC, Surf2Spot delivered better precision–recall trade-offs, as reflected in its AUPRC of 0.601—a 255.6% gain over PPI-hotspotID (0.169) and 670.5% over SPOTONE (0.078) **(Fig. 3a)**. Evaluations based on diverse physicochemical properties **(Fig. 3b)**, ranked Surf2Spot highest,, as did quantitative assessments across the *S*_similarity_, *S*_similarity_, and *SCI* metrics consistently **(Fig. 3c-e)**.

**Fig. 3:**
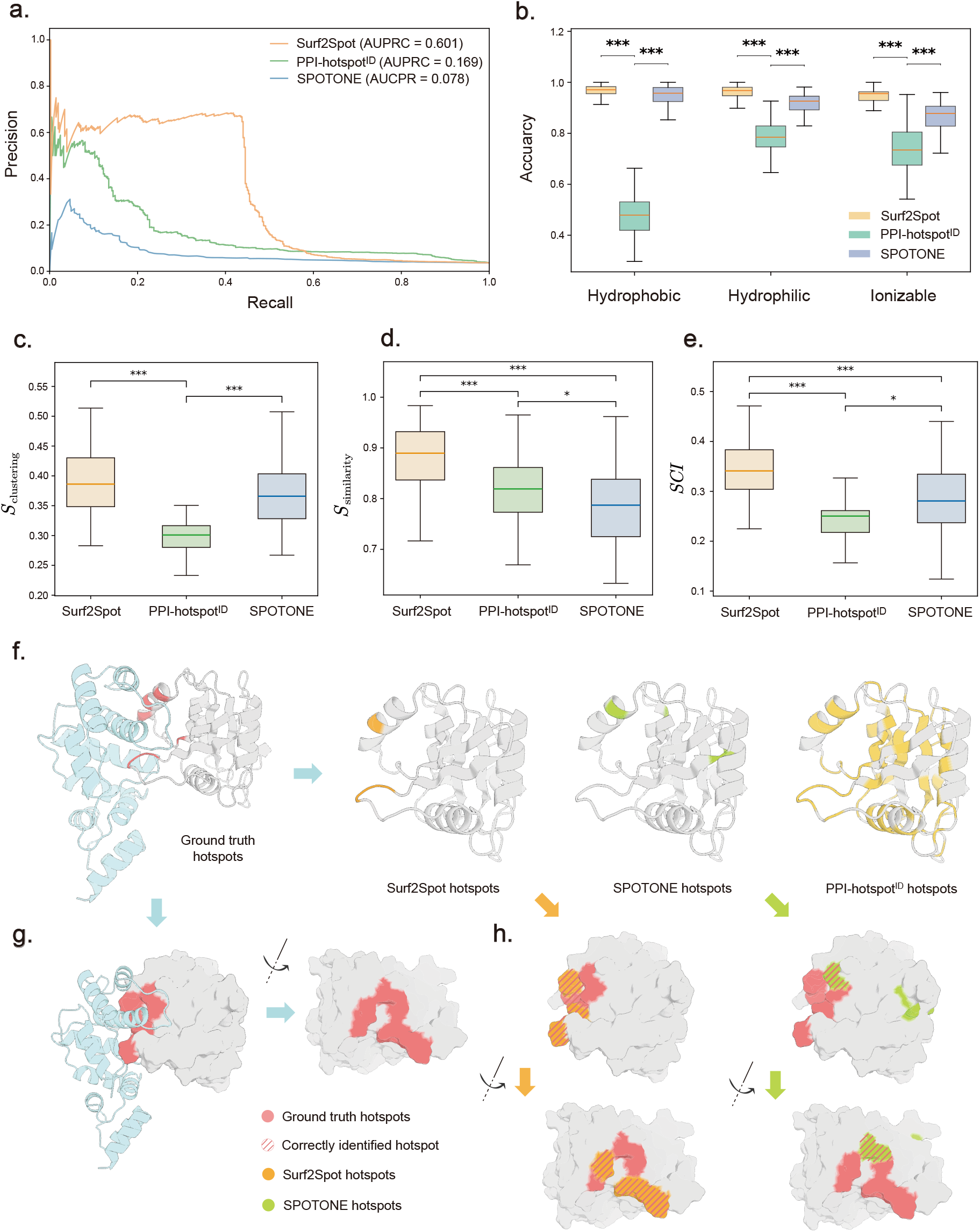
Surf2Spot outperforms state-of-the-art hotspot predictors. **a**, Precision-recall curves of Surf2Spot, PPI-hotspot^ID^, and SPOTONE, with corresponding AUCPR values indicated. **b**, Comparative accuracy of Surf2Spot, PPI-hotspot^ID^, and SPOTONE across different physicochemical amino acid categories. Significance levels: ***p < 0.001, **p < 0.01, *p < 0.05. **c–e**, Quantitative performance comparison of Surf2Spot, PPI-hotspot^ID^, and SPOTONE based on (f)*S*_similarity_, (g) *S*_similarity_, and (h) *SCI* Statistical significance: ***p < 0.001, **p < 0.01, *p < 0.05. **f**, Structure of the *Bos taurus* ADP-ribosylation factor 1 (white) in complex with *Homo sapiens* Arno (blue) (PDB: 1R8S). Experimentally verified hotspots on the lipoprotein are shown in red. **g**, Comparison of hotspot predictions from SPOTONE, PPI-hotspotID, and Surf2Spot against a common ground truth. The ADP-ribosylation factor 1 (white) is shown with its experimental epitope and predicted hotspots (red). **h**, Spatial distribution of predicted hotspots on the protein surface compared with the ground-truth hotspots (red). Predicted residues from SPOTONE (yellow) and Surf2Spot (green) are overlaid for visual comparison.

To demonstrate the predictive performance of Surf2Spot, we conducted a case study using the *Bos taurus* ADP-ribosylation factor 1 (PDB ID: 1R8S, chain A), which forms a complex with *Homo sapiens* Arno **(Fig. 3f)**. Unlike the dispersed residues predicted by SPOTONE or the buried ones identified by PPI-hotspot^ID^, Surf2Spot accurately pinpointed the native interface **(Fig. 3g)**. Surface distribution analysis further confirmed that the Surf2Spot predictions were spatially enriched at the true epitopes **(Fig. 3g)**.

These improvements reflect Surf2Spot’s strength in identifying rare positive residues while minimizing false positives. The flattening tail of its precision–recall curve stems from the inability to assign surface scores to deeply buried residues, which lack accessible surface features and thus are excluded from graph construction. Despite this, Surf2Spot maintains strong classification performance across the full dataset. These results affirm its suitability for real-world applications where hotspot residues are scarce and unevenly distributed.

### Surf2Spot Enhances Binder Design Accuracy and Efficiency

Hotspot residues on a target protein’s surface are critical for guiding structure-based binder design tools such as RFdiffusion^3^, ProteinMPNN^4^, and BindCraft^5^. We hypothesized that Surf2Spot’s predicted hotspots, derived solely from monomeric sequence and structure, could enhance both success rate and computational efficiency, particularly for targets lacking co-crystal structures. To evaluate this, we conducted two case studies: one involving a plant biosynthetic enzyme (NbPDS1)^56^, and the other a fungal virulence factor (VdPDA1)^57^.

NbPDS1 is an enzyme involved in the carotenoid biosynthesis pathway of *Nicotiana benthamiana*^56^ for which no known complex structure has been reported. BLAST alignment identified 5MOG^58^, a homologous pentamer sharing 85% sequence identity, as the closest structural match to NbPDS1 **(Fig. 4a)**. NbPDS1 was divided into three domains (D1–D3), and hotspot residues were independently predicted for each domain (Fig. 4b). Notably, Surf2Spot accurately identified hotspots within a helical region of D3 that mediates oligomerization **(Fig. 4c)**, despite 5MOG being absent from the training set and having <20% similarity to any training sample. This suggests that Surf2Spot effectively captures generalizable physicochemical features of protein interaction interfaces. Based on the predicted hotspots (residues 170, 173, and 174), we selected the D3 segment and performed binder design using both BindCraft and a combined RFdiffusion–ProteinMPNN workflow **(Fig. 4e–f)**. In the BindCraft pipeline, full-length NbPDS1 without hotspot constraints yielded no successful binders after 7.5 days. Restricting design to D3 produced five successful binders in 30.7 hours, and adding Surf2Spot hotspots further reduced design time to 15.6 hours, with a 41% decrease in required iterations **(Fig. 4d, Fig. 4g)**. This demonstrates that hotspot guidance with Surf2Spot improves both efficiency and success rate.

**Fig. 4:**
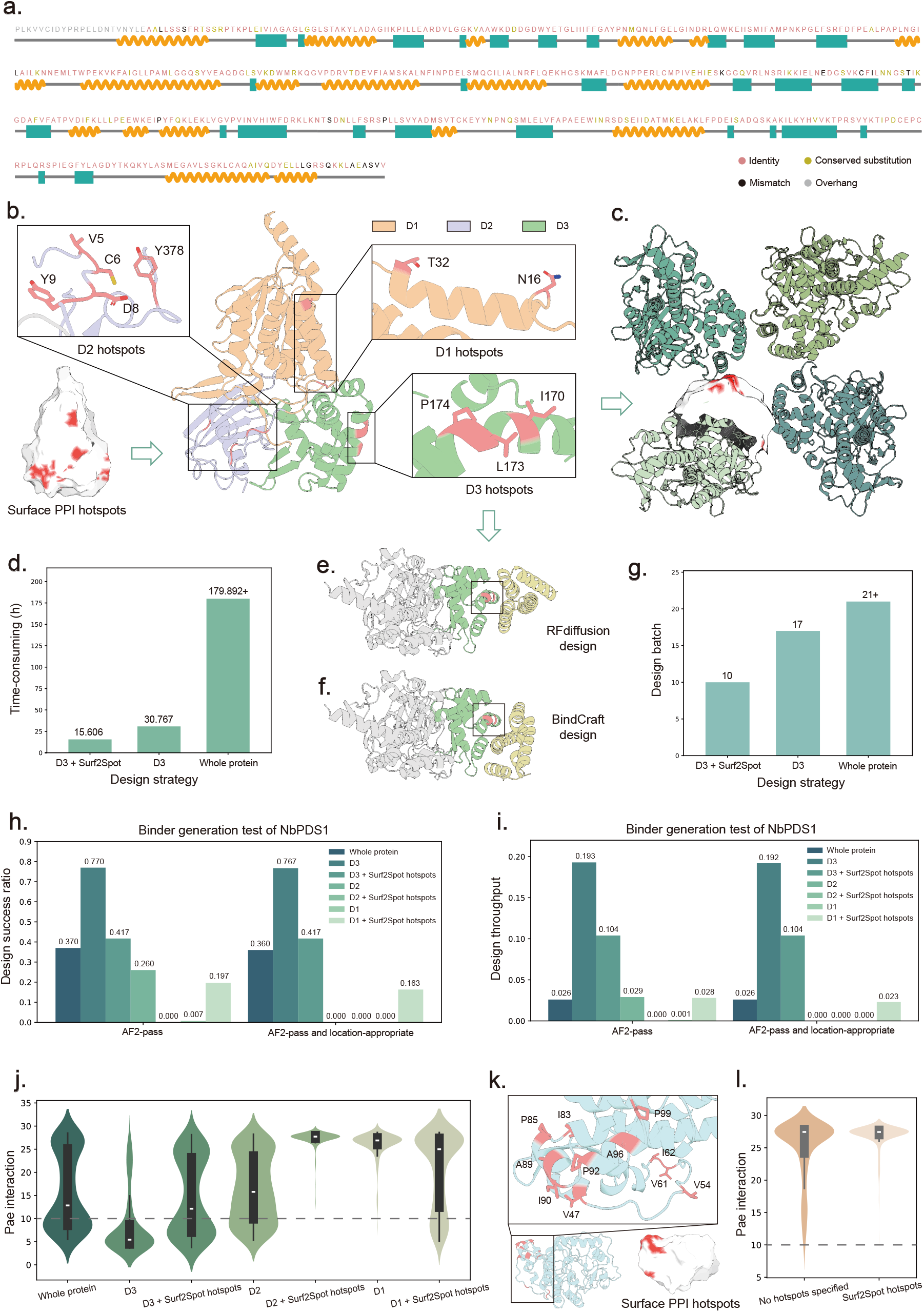
Surf2Spot improves the accuracy and efficiency of binder design. **a**, Architectural features of NbPDS1. Sequence conservation with 5MOG and secondary structure elements are shown along the primary sequence. Identical residues are highlighted in red, conserved substitutions in yellow, and mismatches in grey; overhangs are indicated in black. **b**, Domain-wise prediction of PPI hotspots in NbPDS1. The protein was segmented into three domains—D1 (wheat), D2 (pale green), D3 (blue-white)—with predicted binding hotspots shown for each domain. **c**, Surface-aligned hotspot predictions for 5MOG and the D3 domain of NbPDS1, with predicted hotspots highlighted in red. **e, f**, D3-targeted binder designs for NbPDS1 guided by Surf2Spot-predicted hotspots using RFdiffusion and BindCraft. **d, g**, Computational time and number of attempts required to obtain five successful designs under three strategies: Surf2Spot-guided D3, D3 without hotspot constraints, and full-length NbPDS1. “+” indicates timeout. **h, i**, Success rate and throughput of seven design strategies with or without geometric and AF2-based filtering. **j**, Distribution of interface pAE values among NbPDS1 binders generated under different design strategies. **k**, Structure of VdPDA1 and Surf2Spot-predicted PPI hotspots. The protein backbone is shown in blue-white, with predicted hotspot residues highlighted as red sticks. **l**, Distribution of interface pAE values in binders targeting full-length VdPDA1 with and without Surf2Spot hotspot guidance.

We next benchmarked Surf2Spot-guided designs using RFdiffusion (structure generation) and ProteinMPNN (sequence design), generating 300 candidates per domain on a single NVIDIA A100 GPU. The results revealed strikingly different domain-dependent outcomes. D1 contained two surface hotspots and, although sparse, they enabled 59 viable designs, 49 of which passed all filters. In D2, Surf2Spot localized hotspots predominantly to a disordered region, but all initial and guided designs failed filtering due to structural instability. In D3, the most favorable domain, Surf2Spot produced 231 filtered candidates, including 125 of 125 hotspot-guided designs, with a 65% reduction in generation time (4 vs. 14 minutes) **(Fig. 4h-i; Table S2-1)**. While guidance slightly reduced design diversity, it maintained near-perfect success and yielded the best IPAE scores, with most clustering in the 0–10 range **(Fig. 4j)**. The D3 strategy achieved 0.104 successful designs/min—four times the rate of full-length designs—establishing domain-focused, hotspot-guided design as the preferred paradigm for large proteins **(Table S2-1)**. Together, these findings from the NbPDS1 case study highlight how Surf2Spot-guided hotspot prediction can streamline domain-specific binder design by identifying structurally favorable regions that enhance both success rate and computational efficiency.

By contrast, binder design targeting VdPDA1^57^—a chitin deacetylase secreted by *Verticillium dahliae*—proved highly resistant to binder design. Only 3 of 300 (1%) unguided candidates passed filtering, and none of the 300 hotspot-guided designs (residues 47, 83, 85, 89, 90, 92, 96, 99) were viable (**Fig. 4k-l; Table S2-2**). These results suggest that structural or electrostatic features of VdPDA1 may preclude conventional binder formation. In such cases, alternative scaffolds like nanobodies, which tolerate flexible targets and offer enhanced stability, may represent a more effective design route.

### Surf2Spot Enhances Nanobody Design Efficiency and Interface Quality

We further assessed the utility of Surf2Spot in guiding nanobody design using the IgGM pipeline, targeting the same two proteins, VdPDA1 and NbPDS1. IgGM generated 300 candidate nanobodies within 11.3 h on a single NVIDIA A100 GPU, both with and without incorporating the predicted hotspots. Design success was evaluated using two Rosetta-based performance metrics^59,60^: (1) dG_cross/dSASA × 100, which is inversely correlated with binding strength^61^; and (2) hbonds_int, which represents the number of interface hydrogen bonds and is positively correlated with binding affinity. For VdPDA1—a relatively small protein not subdivided into domains—Surf2Spot predicted three surface hotspots (residues 262, 265, and 268) as likely NAI epitopes **(Fig. 5a)**. With Surf2Spot-guided hotspots, 32.7% of VdPDA1 nanobody designs passed both thresholds, a 3.37-fold increase over the 9.7% success rate without hotspot guidance (**Fig. 5b**). For NbPDS1, Surf2Spot predicted 18 hotspot residues across four NAI epitopes in the D2 and D3 domains (**Fig. 5c**). Guided design yielded higher success rates for three of the four epitopes, with improvements of up to 1.75-fold (**Fig. 5d**).

**Fig. 5:**
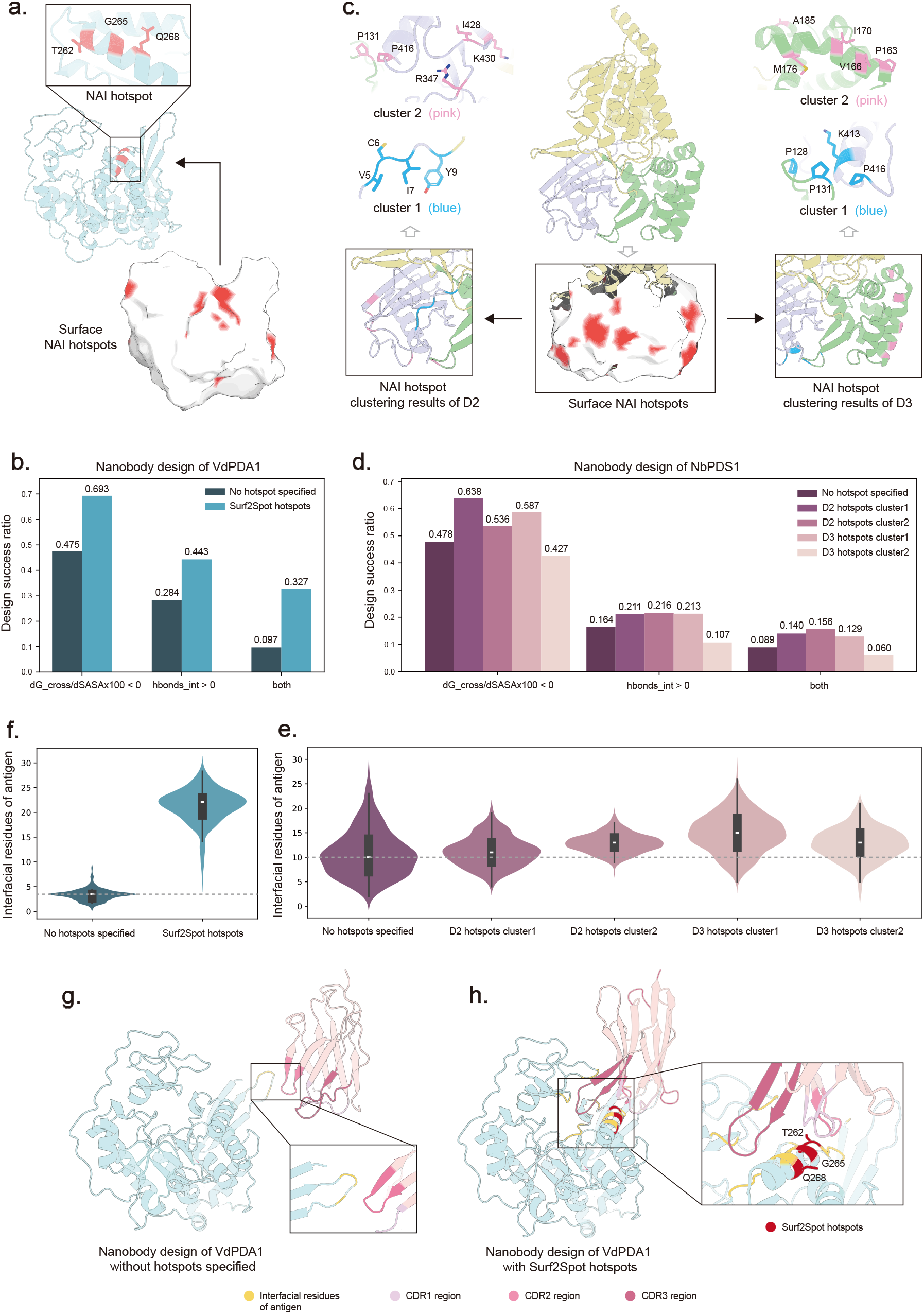
Surf2Spot enhances the accuracy and efficiency of nanobody design. **a**, Structure of VdPDA1 showing NAI hotspots at the surface and residue levels (red). **b, d**, Design success rates and throughput for VdPDA1 and NbPDS1 under three filtering criteria: (1) dG_separated/dSASA × 100, (2) hbonds_int, and (3) both combined. **c**, Domain-level NAI hotspot predictions for NbPDS1 (red), with clustered regions highlighted in blue and pink. **e, f**, Distribution of interfacial residues in (**e**) VdPDA1 binders and (**f**) VdPDA1 nanobodies across different design strategies. The dashed line indicates the peak density for designs without hotspot guidance. **g, h**, Binding poses of nanobody designs generated with and without Surf2Spot hotspot guidance for VdPDA1. Binder interface residues are shown in orange, and hotspots in red.

To further assess binding interface quality, we computed the number of antigen-contacting residues using a ΔSASA > 1.0 Å^2^ threshold. For VdPDA1, Surf2Spot guidance led to a sharp increase in interfacial residue count: guided designs peaked at 25 contacts—nearly five times more than the peak of 4 contacts in unguided designs (**Fig. 5f, Fig. 5g-h**). NbPDS1 designs similarly showed 1.2-to 1.5-fold more interfacial residues when guided by Surf2Spot predictions (**Fig. 5e**). These results demonstrate that Surf2Spot improves not only binder design but also nanobody design, enhancing both success rates and predicted binding interface quality.

## Discussion

Protein binders, engineered molecules that selectively attach to a specific protein target, are foundational tools in structural biology, therapeutics, and synthetic biology. By recognizing precise surface features on a target protein, binders can modulate function, block interactions, recruit effectors, or serve as diagnostic or imaging tools. Their success, however, depends on accurately identifying surface regions that are not only accessible but also energetically favorable for binding. These critical regions, known as interaction hotspots, often contribute disproportionately to binding free energy despite comprising only a small subset of surface residues. Identifying such hotspots remains a significant challenge. Traditional approaches rely on co-crystallized protein complexes, which are expensive and time-consuming to obtain, and typically represent only one of many possible binding modes^62^. Moreover, experimental methods like alanine scanning mutagenesis are labor-intensive and low-throughput. Computational tools for hotspot prediction have advanced in recent years, but many, such as HotPoint^14^, KFC2^15^, and PredHS2^17^ rely on complex structural data and energy-based modeling, which limits their applicability to proteins with known interaction partners. Others, like SPOTONE, use sequence-derived information from multiple sequence alignments, further restricting their use in novel or poorly characterized proteins. These constraints limit their utility for binder design against novel or structurally uncharacterized proteins.

Surf2Spot was developed to address this gap. By leveraging predicted structures and surface-based graph representations, it enables accurate hotspot prediction directly from a single protein sequence, without requiring co-crystal structures, multiple sequence alignments, or known binding partners. This architecture dramatically reduces both time and cost compared to traditional methods such as alanine scanning or crystallographic structure determination, which typically require months of experimental work. Surf2Spot can generate high-confidence hotspot predictions in minutes, making it well-suited for high-throughput screening and rapid iteration in protein engineering workflows. By combining surface-aware modeling with pretrained sequence and structure embeddings, Surf2Spot unlocks scalable, partner-agnostic hotspot discovery, even for structurally uncharacterized or “hard-to-drug” targets^63^.

Surf2Spot’s design addresses key technical challenges that have historically limited hotspot prediction tools. One of the central difficulties is the severe label imbalance in available datasets: true interaction hotspots are sparse, and experimentally validated complexes capture only a subset of potential interactions for any given protein. To mitigate this, Surf2Spot adopts a multi-pronged strategy. First, it employs surface-focused representations that exclude buried residues unlikely to participate in protein–protein interactions. Second, it decomposes proteins into functional substructures using GPSite and Chainsaw, narrowing the prediction space to regions more likely to be involved in binding. Finally, it uses a focal loss function during training to emphasize hard-to-classify examples, improving model performance under class imbalance^64^. Together, these innovations allow Surf2Spot to outperform state-of-the-art sequence-based and structure-based methods across multiple evaluation metrics, including F1 score, Matthews correlation coefficient, and AUPRC.

Beyond predictive accuracy, Surf2Spot’s practical value lies in its ability to guide and streamline protein design. In binder engineering workflows using tools like RFdiffusion^3^, ProteinMPNN^4^, and BindCraft^5^, hotspot constraints provided by Surf2Spot substantially reduced design time and improved success rates. In the NbPDS1 case study, hotspot guidance cut computational time by half and increased the rate of viable candidates fourfold. Similarly, Surf2Spot-guided nanobody design using IgGM^6^ resulted in significantly higher interface quality and a greater number of predicted contact residues, particularly for challenging targets like VdPDA1. These improvements demonstrate Surf2Spot’s utility in focusing design efforts on structurally favorable regions, avoiding disordered or inaccessible sites, and enabling more efficient iteration.

Importantly, Surf2Spot’s hotspot predictions also retain physicochemical interpretability, as they are derived from geometric and residue-level features of the protein surface. This allows researchers to rationally inspect and refine candidate designs based on known principles of interface energetics. Compared to PPI-hotspot^ID^, which emphasizes residues contributing to complex-wide stability, Surf2Spot is optimized to identify anchor residues that initiate and stabilize direct interactions. Similarly, tools like SPOTONE^18^ and KFC2^15^ rely heavily on multiple sequence alignments or known complex structures, limiting their use for proteins lacking homologs or partner data. In contrast, Surf2Spot operates solely on predicted monomeric structures, enabling broader applicability in orphan or non-model proteins. This focus yields more actionable predictions for structure-based design, as confirmed by our benchmarking results.

While Surf2Spot was developed and validated using general protein datasets, its application to plant proteins in this study highlights an important emerging use case. In many non-model systems, including plants, fungi, and pathogens, protein interaction data and structural resources are limited. Surf2Spot’s ability to operate on predicted monomeric structures makes it particularly suitable for binder development in these contexts. Its success in guiding binder and nanobody design for plant enzymes and pathogen effectors suggests broader potential in agricultural biotechnology, such as engineering disease resistance or modulating key metabolic pathways. Moreover, in emerging applications such as bioPROTACs, where a binder is used to recruit target proteins to E3 ligases, the ability to design high-affinity, non-antibody binders or nanobodies from minimal structural information will be increasingly valuable.

In conclusion, Surf2Spot offers a scalable, accurate, and interpretable approach to hotspot prediction that enables more efficient and effective protein binder design. By operating directly on predicted structures and bypassing the need for experimental complex data, it expands the range of tractable targets and accelerates the design-build-test cycle across diverse applications in biology, biotechnology, and synthetic design.

## Methods

### Surface feature representation

In our framework, a protein surface represented as a point cloud is modeled as an undirected graph *G=(V,E)*, where *V* denotes the surface points (nodes) and *E* denotes the contacts between these points derived from the triangular mesh (edges). Thus, the surface graph *G* can be represented by a node feature matrix *X* and an adjacency matrix *A* To train our model, we extracted three groups of protein correlation features: sequence properties (ProtTrans embedding and AA type), structural properties (DSSP), and surface properties (point cloud sampling), which were concatenated after processing by MLPs to form the final node feature matrix.

#### Sequence properties

Sequence embeddings were computed using the pre-trained protein language model ProtTrans, which was trained on 393 billion amino acids from UniRef and BFD^65,66^. Each amino acid was encoded as a 1024-dimensional vector using ProtTrans with default parameters. In parallel, amino acid types were encoded using one-hot encoding.

#### Structural properties

Three types of structural properties were computed using the DSSP program: (i) a 9-dimensional one-hot encoded secondary structure profile, where the first 8 dimensions represent 8 categories of secondary structure states and the last dimension represents unknown secondary structure; (ii) peptide backbone torsion angles PHI and PSI; and (iii) solvent-accessible surface area (ASA), which was normalized to relative solvent accessibility (RSA) using the maximal possible ASA of the corresponding amino acid type.

#### Surface properties

One geometric and three energetic features were extracted and attached to the nodes of the pocket surface. The geometric feature included the shape index; the energetic features included Poisson-Boltzmann continuum electrostatics, hydrogen bond potential, and hydropathy.

#### Shape Index

The shape index describes local surface topology in terms of principal curvature and was proposed by Koenderink and van Doorn as a single-valued angular measure. The definition is as follows:

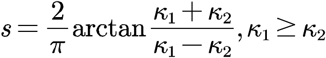

where κ_1_ and κ_2_ are the principal curvatures of the surface at a specific point, which can be calculated from Gaussian curvature and mean curvature as follows:

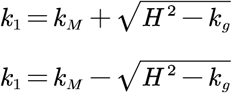

#### Poisson-Boltzmann continuum electrostatics

The Poisson-Boltzmann equation describes the continuous charge distribution of biomolecules in solution as follows:

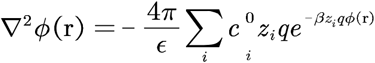

Where *ϕ* is the system potential; *ϵ* is the dielectric constant in solution;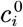 and *z*_*i*_ are the bulk phase concentration and charge of the i-th atom, respectively; *β=*1*/k*_*B*_*T*, where *k*_*B*_ is the Boltzmann constant. The calculation was performed by PDB2PQR and APBS following the MaSIF method.

#### Hydrogen-bond potential

A hydrogen bond is essentially a dipole-dipole interaction, usually represented as X-H…Y, where X is covalently (or ionically) bonded to hydrogen and has high electronegativity, while Y is generally an atom containing lone pairs of electrons. Hydrogen bonding interactions are widely used in the construction of force fields. Similar to previous work, hydrogen-bond potential was assigned to surface nodes.

#### Hydropathy

Hydrophobic interaction is a non-covalent interaction present between non-polar groups and is prevalent in biological systems. Similar to previous work, hydrophobic interactions were assigned to surface nodes based on the residues they belonged to.

### Compilation of training datasets

The benchmark dataset for evaluating PPI-binding hotspot prediction was derived from the supplementary materials of the PDBBIND2020 and GPSite datasets^67,38^. The benchmark dataset for predicting NAI-binding hotspots was derived from the VHH antigen-antibody conjugate structures in SabDab^54^. A total of 1200 protein dimer complexes corresponding to target proteins were collected from PDBBIND2020, and 446 were supplemented from the GPSite dataset. SabDab included 1248 antigen VHH antibody complex structures corresponding to antigens. We removed redundant proteins with sequence length differences greater than 20% and shared sequence identities greater than 80%, and removed proteins shorter than 30 residues. A binding residue was defined if the ΔSASA between the target residue and ligand was greater than 1.0 and the difference in binding energy between FoldX alanine mutation scanning was greater than the specified threshold, which is 2 kcal/mol for PPI and 0.01kcal/mol for NAI^68^. Complex structures with fewer than three hotspot residues were removed to ensure the quality of the dataset. To prevent the model from over-fitting the amino acid conformation of the binding state on the contact surface of the complex, ESMFold was used to predict the monomer conformation of the target protein in the complex and supplement it into the dataset. The final PPI dataset contains 1020 proteins, and the NAI dataset contains 498 proteins. Each pair of complex structure and its corresponding monomer prediction was assigned to either the training set or the test set together, ensuring consistency and fairness in the evaluation process.

### Compilation of model evaluation datasets (Test_52)

We adopted three publicly available and widely used benchmark datasets from previous studies: Dset_72^69^, Dset_164 and Dset_186^70^—named according to the number of proteins they contain. We selected protein structures from these established benchmark datasets that were absent from our training data. Initial screening identified 11, 24, and 50 candidate proteins from Dset_72, Dset_186, and Dset_164, respectively. We processed the dataset by removing duplicate sequences using the same method applied to the training dataset and required each entry to contain at least six hotspot residues. These candidates were further refined using ESMFold to retain only high-confidence monomeric structures with pLDDT scores >80, yielding a final fused test set (Test_52) comprising 10 proteins from Dset_72, 18 from Dset_186, and 24 from Dset_164. This rigorous selection process ensured that Test_52 represented a diverse, non-redundant collection of structurally reliable targets for robust model validation.

### Quantifying the spatial distribution of hotspots

To evaluate the spatial organization and predictive accuracy of hotspot residues, we introduced several quantitative metrics that assess how well the predicted hotspots capture key structural and distributional characteristics. These include the Spatial Consensus Index (*SCI*), which measures the consistency of hotspot locations across predictions; the clustering score (*S*_clustering_), which quantifies spatial compactness and uniformity; and the similarity score (*S*_similarity_), which measures the spatial overlap between predicted hotspots and reference hotspots.

*SCI* reflects the degree of consistency in hotspot spatial distribution and quantifies the comprehensive quality of hotspot distribution and is defined as:

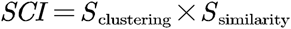

The clustering score *S* _clustering_ is defined as the product of the *U*_hotspots_ and *C*_hotsopts_, which requires hotspots to be both compact and evenly distributed:

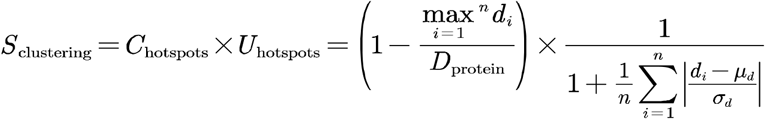

*U*_hotspots_ measures the uniformity of hotspot distribution, with values closer to 1 indicating a more uniform distribution (i.e., all points are at similar distances from the center). In contrast, lower values indicate a less uniform distribution, where some points are very close to the center while others are much farther away:

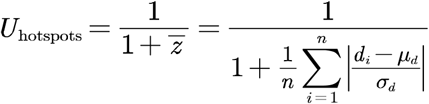

**r**_*i*_: CA atom coordinate vector of the i-th hotspot residue

**c**: Geometric center of the hotspot residue

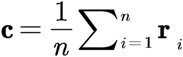

**a**_*j*_: Coordinate vector of the j-th atom in the protein

*n*: Number of hotspot residues

*m*: Number of protein atoms Distance calculation:

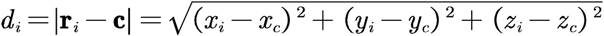

Statistical calculation

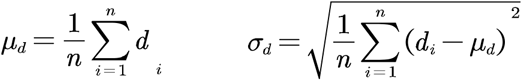

*C*_hotspots_ measures the spatial compactness of hotspots within a protein. A value closer to 1 indicates that the hotspots are more tightly clustered, whereas a smaller value suggests a more dispersed spatial distribution:

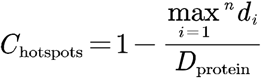

*D*_protein_: maximum diameter of the protein

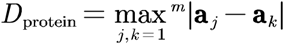

The similarity score *S*_similarity_ quantifies the proximity between the geometric centers of two hotspot residue sets in three-dimensional space. A value close to 1 indicates that the two hotspot centers are nearly coincident, reflecting high prediction accuracy, whereas a value close to 0 indicates that the centers are far apart, imp;lying low prediction accuracy. Standardization ensures comparability of the scores among proteins of different sizes:

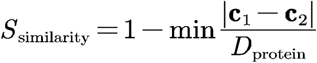

### Construction and training of DGCNN and GAT networks

#### Model of DGCNN

We computed a directed graph *G=(V,E)* representing the local point cloud structure, where *X*={*x*_1_,…,*x*_*n*_} ⊆R^*F*^ and *Ē* ⊆*V*×*V* are the vertices and edges, respectively. In the simplest case, we constructed *G* as the k-nearest neighbor (*k*-NN) graph of X in *R*^*F*^. The graph includes a self-loop, meaning each node also points to itself. We defined edge features as *e*_*ij*_=*h*_Θ_(*x*_*i*_,*x*_*j*_), where *h*_Θ_:*R*^*F*^ × *R*^*F*^ → *R*^*F*′^ is a nonlinear function with a set of learnable parameters Θ.

Then, a channel-wise symmetric aggregation operation □ (e.g., sum or max) is applied to re-characterize the central point, aggregating additional neighborhood information. Overall, given an *F*-dimensional point cloud with n points, EdgeConv produces an *F*^′^-dimensional point cloud with the same number of points:

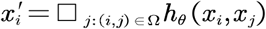

It has been shown that recomputing the graph using nearest neighbors in the feature space produced by each layer is beneficial, which is a crucial distinction between Dynamic Graph CNNs (DGCNNs) and graph CNNs that operate on a fixed input graph.

#### Construction of models

The above-mentioned attributed graph, incorporating features from ProtTrans, DSSP, and surface properties, is input into several MLP layers. The processed surface is represented by *G*=(*V,E*), where *X*={*x*_1_,…,*x*_*n*_} ⊆R^*F*^ and *Ē* ⊆*V*×*V* are the vertices and edges, respectively. We constructed *G* as the k-nearest neighbor (*k*-NN, k=20) graph of X in.*R*^*F*^ The graph includes a self-loop, meaning each node also connects to itself. Edge features are defined as *e*_*ij*_=*h*_Θ_(*x*_*i*_,*x*_*j*_),, where *h*_Θ_:*R*^*F*^ × *R*^*F*^ → *R*^*F*′^ is a nonlinear function with a set of learnable parameters. Θ The new representations are aggregated and input into a three-layer DGCNN. Each DGCNN layer includes message passing, edge update, and global node update modules to learn residue representations by considering multi-scale interactions at the node, edge, and global context levels. DGCNN applies a channel-wise symmetric aggregation operation to re-characterize each central point and recomputes the graph using nearest neighbors in the feature space produced by each layer. Overall, given an F′-dimensional point cloud with n points, EdgeConv produces an F’-dimensional point cloud with the same number of points. In this way, the three-layer DGCNN progressively updates the structural edge representations of the protein surface map and aggregates node features.

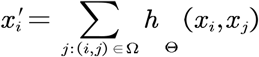

The node features output by DGCNN are combined with the surface representations via a residual connection before being input into the graph neural network module. For NAI hotspot prediction, the representation after this residual connection is fed into a 3-layer Graph Attention Network (GAT), and its output is then combined with the DGCNN output through a second residual connection. The output of the last graph layer is passed through an MLP to predict the protein interaction probabilities for all amino acid residues:

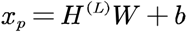

Where *H*^(*L*)^ is the output of the Lth graph neural network layer;*W* is the weight matrix;*b* is the bias term, and *x*_*P*_ is the prediction of the surface node. The discrimination of the output signal *x*_*P*_ is enhanced by introducing a magnification *α* factor of 5 times, and finally the binding probability value *P*_*P*_ of each surface point is output using the Sigmoid activation function.

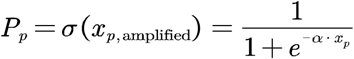

#### Training process

We performed five-fold cross-validation on the training data, where in each iteration, the model was trained on four folds and evaluated on the remaining fold. This process was repeated five times, and the average validation performance was used to optimize the network’s hyperparameters. In the test phase, all five trained models from cross-validation were used to make predictions, which were then averaged to produce the final Surf2Spot output. The Adam optimizer (learning rate = 1e-4) was used for model optimization on the focal loss:

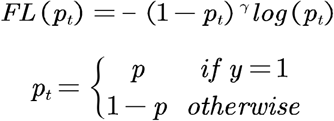

The training process (batch_size=1) lasted at most 120 epochs, and we performed early stopping based on validation performance, which took ∼18 hr on an NVIDIA A100 GPU.

### Model performance evaluation

We used accuracy (ACC), precision, recall, F1-score (F1), Matthews correlation coefficient (MCC), area under the receiver operating characteristic curve (AUROC), and area under the precision-recall curve (AUPRC) to measure predictive performance:

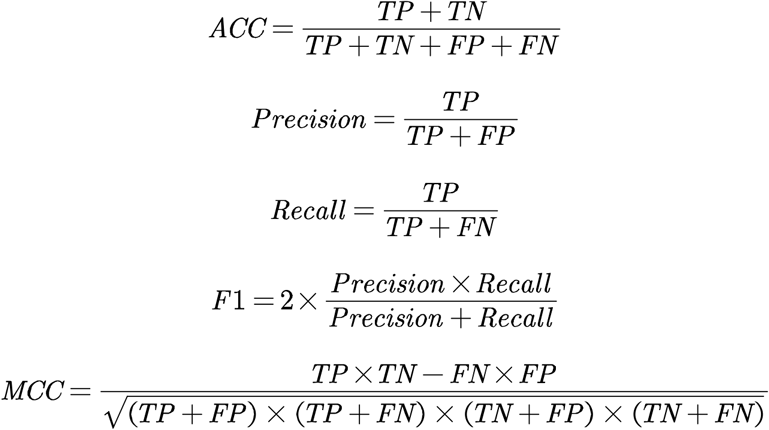

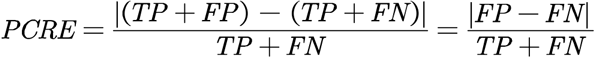

PCRE (Predicted Count Relative Error) is defined as the absolute value of the difference between the number of predicted positive samples and the number of true positive samples, divided by the number of true positive samples. This metric reflects the relative error in the count of positive samples predicted by the model.

True positives (TP) and true negatives (TN) denote the number of interacting and non-interacting sites identified correctly, and false positives (FP) and false negatives (FN) denote the number of incorrectly predicted interacting and non-interacting sites, respectively. AUROC and AUPRC are independent of thresholds, thus revealing the overall performance of a model. Other metrics were calculated using a threshold to convert predicted interaction probabilities to binary predictions, which was determined by maximizing the F1-score for each model.

Our model defines a hotspot amino acid based on the principle of spatial proximity: any residue is identified as a hot spot if its 10-angstrom spatial sphere contains surface points predicted as hot spots. This rule effectively mitigates identification biases stemming from point cloud mapping variations, aligning predictions more closely with the authentic spatial patterns of biophysical interactions.

### Binder design pipeline and filtration criteria

#### RFdiffusion design comparison

To compare the design efficiency of RFdiffusion with or without Surf2Spot, we designed binders to two targets: NbPDS1 (AlphaFoldDB ID: AF-B1NYI4-F1-v4) and VdPDA1 (AlphaFoldDB ID: AF-G2XC45-F1-v4). In the case of Surf2Spot guidance, we targeted binders to specific sites on the target protein using predicted hotspot residues. The selected hotspots and truncated substructures are listed below (with chain and residue indices from PDB); all other parameters were kept at their default values.

We set the length of the generated binders to 100 amino acids and generated 300 binder structures per run, with all other parameters left as defaults. For sequence design and refinement, we applied the previously described ProteinMPNN-FastRelax protocol^71^, which starts with an initial ProteinMPNN round and then iteratively alternates between FastRelax and ProteinMPNN to progressively optimize sequence– structure agreement. One binder sequence was generated for each structure produced by RFdiffusion. To filter the designs, we ran AlphaFold2 with initial guess and target templating. A design was classified as successful if it met two criteria: (1) the predicted aligned error (pAE) of interaction between the binder and target < 10, and the predicted local distance difference test (pLDDT) > 70. All AlphaFold2 parameters were left at their default settings.

#### BindCraft design comparison

To compare the design efficiency of BindCraft with or without the guidance of Surf2Spot, we designed the same target proteins as in the efficiency comparison experiment of RFdiffusion. Under the guidance of Surf2Spot, we used the predicted hotspot residues to direct the conjugate to a specific site on the target protein. The selected hotspots and truncated substructures are as follows (chain and residue indices from PDB):

For the BindCraft design, the length of the adhesive was fixed at 100 amino acids, and the expected number of final filtered designs was set to 5. All other design and filtering parameters were kept at the default settings.

Default filtering parameters that were used include:

a. AF2 confidence pLDDT score of the predicted complex (> 0.8)
b. AF2 interface predicted confidence score (i_pTM) (> 0.5)
c. AF2 interface predicted alignment error (i_pAE) (> 0.35)
d. Rosetta interface shape complementarity (> 0.55)
e. Number of unsaturated hydrogen bonds at the interface (< 3)
f. Hydrophobicity of binder surface (< 35%)
g. RMSD of binder predicted in bound and unbound form (< 3.5 Å)

### Nanobody design pipeline and filtration criteria

To evaluate the impact of Surf2Spot on the design efficiency of IgGM, we generated nanobodies targeting two distinct antigens: NbPDS1 (AlphaFoldDB ID: AF-B1NYI4-F1-v4) and VdPDA1 (AlphaFoldDB ID: AF-G2XC45-F1-v4). In the Surf2Spot-assisted design group, nanobodies were strategically engineered to bind predicted hotspot residues within specific antigenic epitopes. The selected hotspots and their corresponding truncated structural domains were annotated based on chain identifiers and residue indices from reference PDB structures. The selected hotspots and truncated substructures are listed below (chain and residue indices are based on the PDB entry); all other parameters were kept at their default settings.

#### Nanobody binding free energy and interaction analysis

The binding energy and interface properties of nanobody-antigen complexes were analyzed using Rosetta (v3.13) through the following steps: (1) E

##### nergy minimization

The minimize.static.linuxgccrelease module was executed with parameters -min_type lbfgs_armijo_nonmonotone (L-BFGS optimization), -score:weights ref2015_cst (ref2015_cst scoring function), -min_tolerance 0.001 (convergence threshold), and backbone constraints (-relax:bb_move false). (2) Interface analysis: The InterfaceAnalyzer.static.linuxgccrelease tool quantified interface metrics, specifying the interaction chains (-interface A_H), enabling repacking of separated states (-pack_separated), and outputting energy/structural features.

#### Nanobody design filtering criteria

Key parameters were extracted and filtered based on: (1) Binding efficiency: Complexes with dG_cross/dSASA × 100 < 0 were retained, indicating favorable binding free energy (dG_cross) per unit solvent-accessible surface area (dSASA). (2) Hydrogen bonds: On the basis of ensuring dG_cross/dSASA × 100 < 0, interfaces with hbonds_int > 0 were preferred to ensure specific polar interactions. Results were sorted by dG_cross/dSASA × 100 (ascending) and exported as a CSV file, including comprehensive metrics (total score, dG_cross/dSASA × 100, hbonds_int, total_score, etc.).

## Data availability

The predicted structures of NbPDS1 and VdPDA1 can be accessed at the AlphaFold Protein Structure Database under accession codes <u>B1NYI4</u> and <u>G2XC45</u>, respectively. Protein structures from the PDB used for training data construction are available at https://huggingface.co/datasets/anwzhao/Surf2Spot_raw_data.

## Code availability

The Surf2Spot source code is freely available for academic use at https://github.com/AnwZhao/Surf2Spot.

## Acknowledgement

This research was supported by multiple funding sources. We gratefully acknowledge the financial support from the Beijing Rural Revitalization Agricultural Science and Technology Project (NY2601491026) and the Major Science and Technology Special Project of Liaoning Province (2025JH1/11700012). Additional support was provided by the Guiding Special Fund for Central Universities to Build World-Class Universities (Disciplines) and Promote Characteristic Development (2025AC030) and the Pinduoduo-China Agricultural University Research Fund (PC2024A02002).

## Supplementary table S1 to S4

**Table S1-1:** Test_52 global amino acid evaluation.

| Model | Basis of prediction | ACC | Precision | Recall | F1 | MCC | AUROC | AUPRC | PCRE |
| --- | --- | --- | --- | --- | --- | --- | --- | --- | --- |
| Surf2Spot | surface | 0.982 | 0.611 | 0.609 | 0.610 | 0.602 | 0.800 | 0.601 | 0.008 |
| PPI-hotspot<sup>ID</sup> | structure | 0.630 | 0.077 | 0.836 | 0.141 | 0.175 | 0.774 | 0.169 | 9.902 |
| SPOTONE | sequence | 0.942 | 0.162 | 0.142 | 0.151 | 0.122 | 0.617 | 0.078 | 0.127 |

**Table S1-2:** Test_52 protein average evaluation.

| Model | Basis of prediction | ACC | Precision | Recall | F1 | MCC | AUROC | AUPRC | PCRE |
| --- | --- | --- | --- | --- | --- | --- | --- | --- | --- |
| Surf2Spot | surface | 0.977 | 0.618 | 0.586 | 0.562 | 0.571 | 0.788 | 0.602 | 0.474 |
| PPI-hotspot<sup>ID</sup> | structure | 0.654 | 0.106 | 0.832 | 0.181 | 0.205 | 0.785 | 0.239 | 9.874 |
| SPOTONE | sequence | 0.932 | 0.211 | 0.142 | 0.150 | 0.130 | 0.599 | 0.130 | 0.532 |

**Table 2-1:**
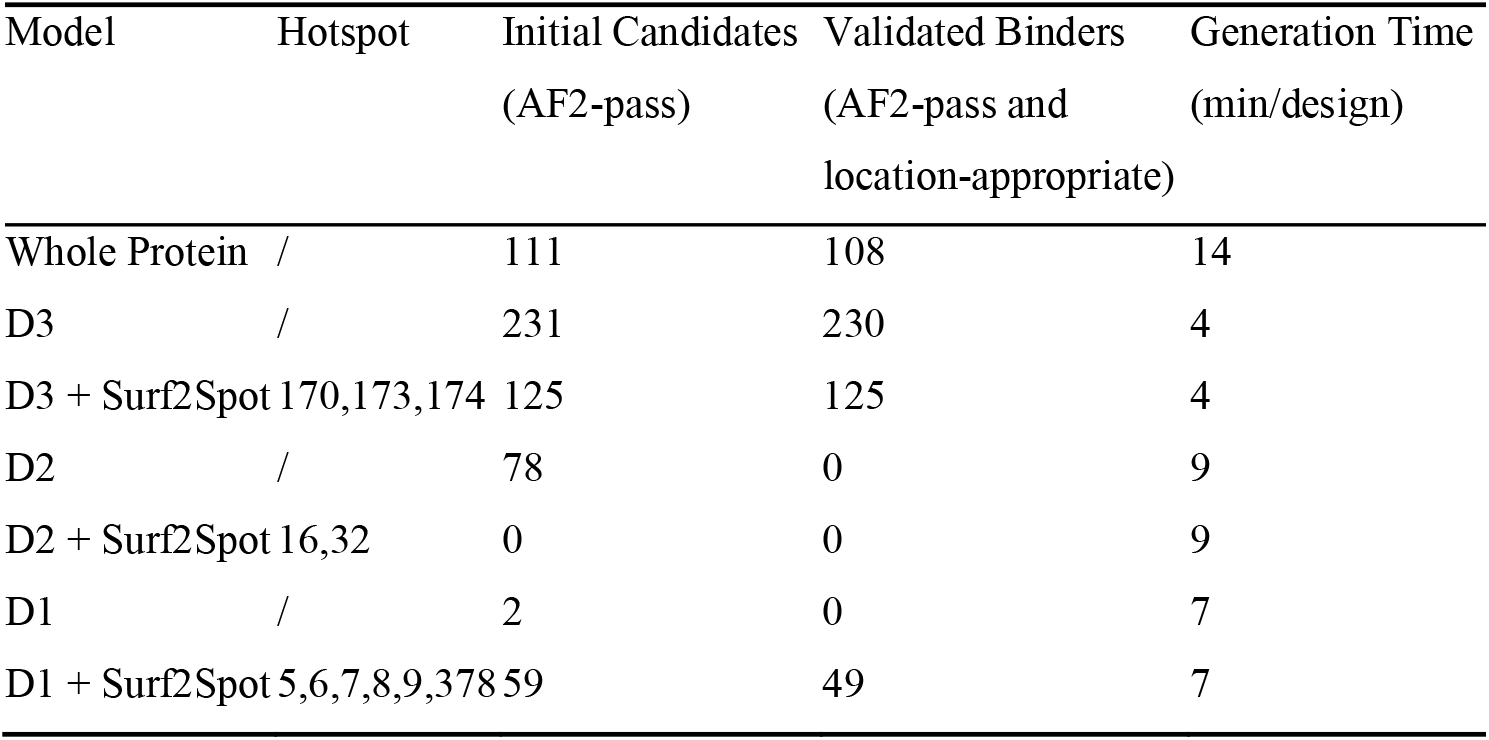
*Nb*PDS1 hotspot evaluation.

**Table S2-2:** VdPDA1 hotspot evaluation.

| Model | Hotspot | Initial Candidates<br>(AF2-pass) | Validated Binders<br>(AF2+position) | Generation Time<br>(min/design) |
| --- | --- | --- | --- | --- |
| Whole Protein | / | 3 | 3 | 14 |
| Whole Protein + Surf2Spot | 47,83,85,89, 90,92,96,99 | 0 | 0 | 14 |

**Table S3-1:** RFdiffusion binder design parameter of NbPDS1.

| Design Strategy | Amino Acid Range | Hotspots |
| --- | --- | --- |
| Whole Protein | 1-586 | / |
| D3 | 136-238 | / |
| D3 + Surf2Spot | 136-238 | 170, 173, 174 |
| D2 | 1-32/75-158/238-273/334-441 | / |
| D2 + Surf2Spot | 1-32/75-158/238-273/334-441 | 5, 6, 7, 8, 9, 378 |
| D1 | 11-115/250-333/445-508 | / |
| D1 + Surf2Spot | 11-115/250-333/445-508 | 16, 32 |

**Table S3-2:** RFdiffusion binder design parameter of VdPDA1.

| Design Strategy | Amino Acid Range | Hotspots |
| --- | --- | --- |
| Whole Protein | 1-586 | / |
| Whole Protein | 1-586 | 47, 85, 89, 90, 92, 96, 99 |

**Table S3-3:** BindCraft binder design parameter of NbPDS1.

| Design Strategy | Amino Acid Range | Hotspots |
| --- | --- | --- |
| Whole Protein | 1-586 | / |
| D3 | 136-238 | / |
| D3 + Surf2Spot | 136-238 | 170, 173, 174 |

**Table S3-4:** BindCraft binder design parameter of VdPDA1.

| Design Strategy | Amino Acid Range | Hotspots |
| --- | --- | --- |
| Whole Protein | 1-586 | / |
| Whole Protein | 1-586 | 47, 85, 89, 90, 92, 96, 99 |

**Table S4-1:** IgGM nanobody design parameter of NbPDS1.

| Design Strategy | Hotspots |
| --- | --- |
| No hotspot | / |
| hotspot cluster1 on domain1 | 131, 347, 416, 428, 430 |
| hotspot cluster2 on domain1 | 5, 6, 7, 9 |
| hotspot cluster1 on domain2 | 163, 166, 170, 176, 185 |
| hotspot cluster2 on domain2 | 128, 131, 413, 416 |

**Table S4-2:** IgGM nanobody design parameter of VdPDA1.

| Design Strategy | Hotspots |
| --- | --- |
| No hotspot | / |
| Surf2Spot hotspot | 262, 265, 268 |

**Supplementary Fig. 1**.

**a and b**, B-factor distributions (bound states) and pLDDT scores (unbound states) for the PPI and NAI datasets.

**Supplementary Fig. 2**.

**a–g**, Accuracy, precision, recall, AUROC, AUCPR, F1-score, and MCC at the global amino acid level across Surf2Spot, PPI-hotspot^ID^, and SPOTONE (light blue). Corresponding comparisons at the protein-average level are shown in dark blue.

**Supplementary Fig. 3**.

**a and b**, Adjacency matrices of VdPDA1 and NbPDS1.

**Supplementary Fig. 4**.

**a and b**, Domain architecture and PPI site enrichment of NbPDS1. **c and d**, Domain architecture and PPI site enrichment of VdPDA1.

