## Supplementary figures and images for "Surf2Spot: A Surface-Informed Geometry-Aware Model for Predicting Binder and Nanobody Design Hotspots"

### Supplemental Figure 1

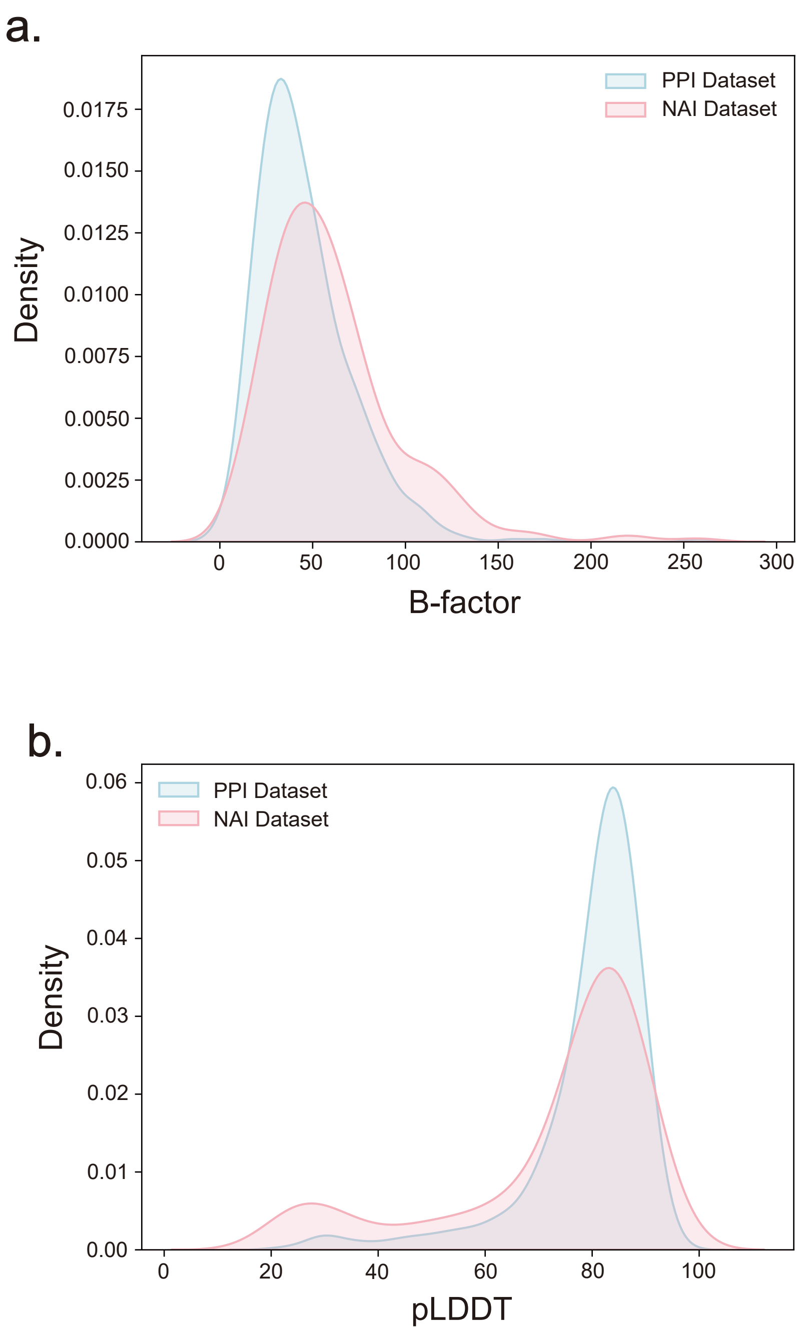

### Supplemental Figure 2

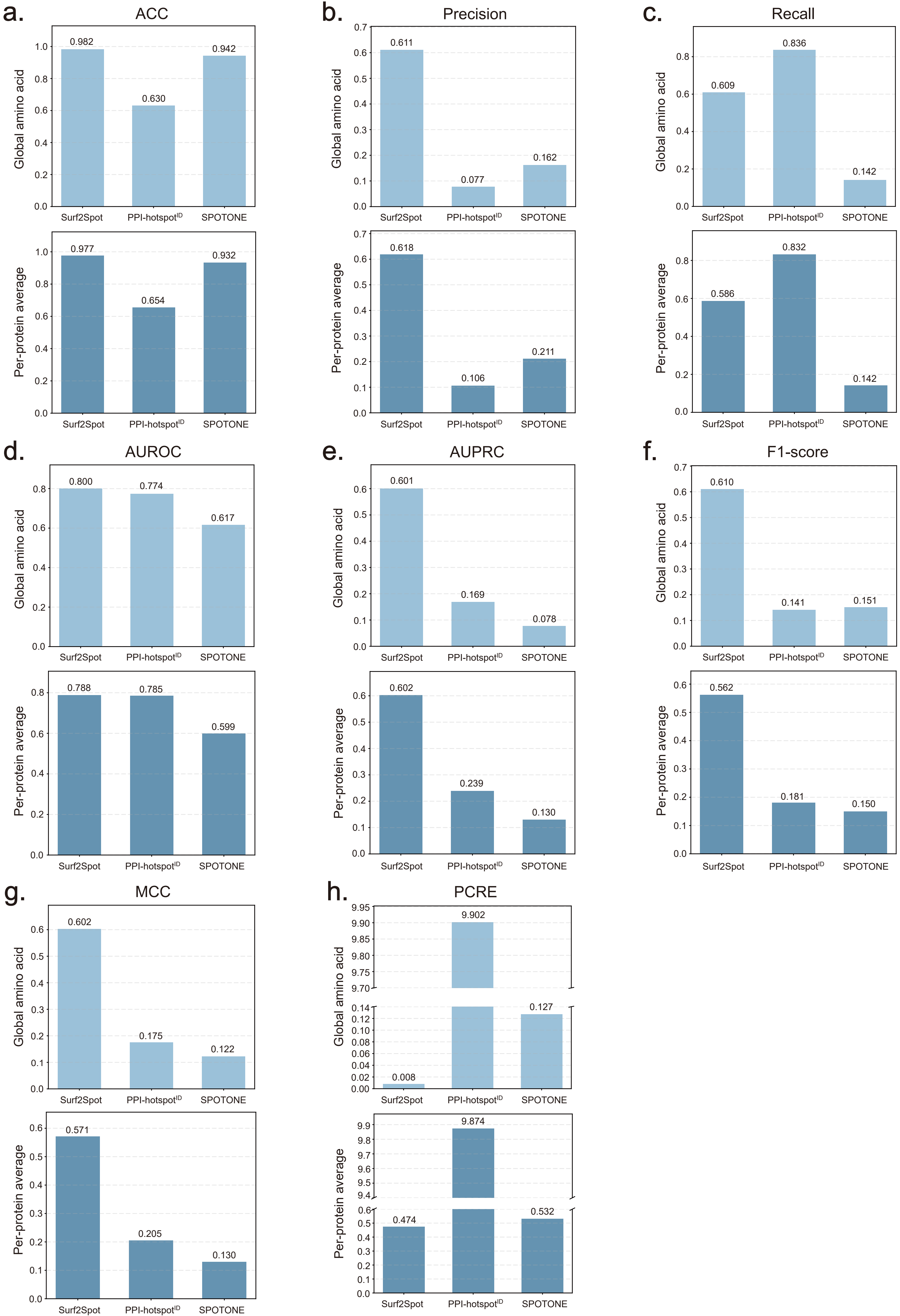

### Supplemental Figure 3

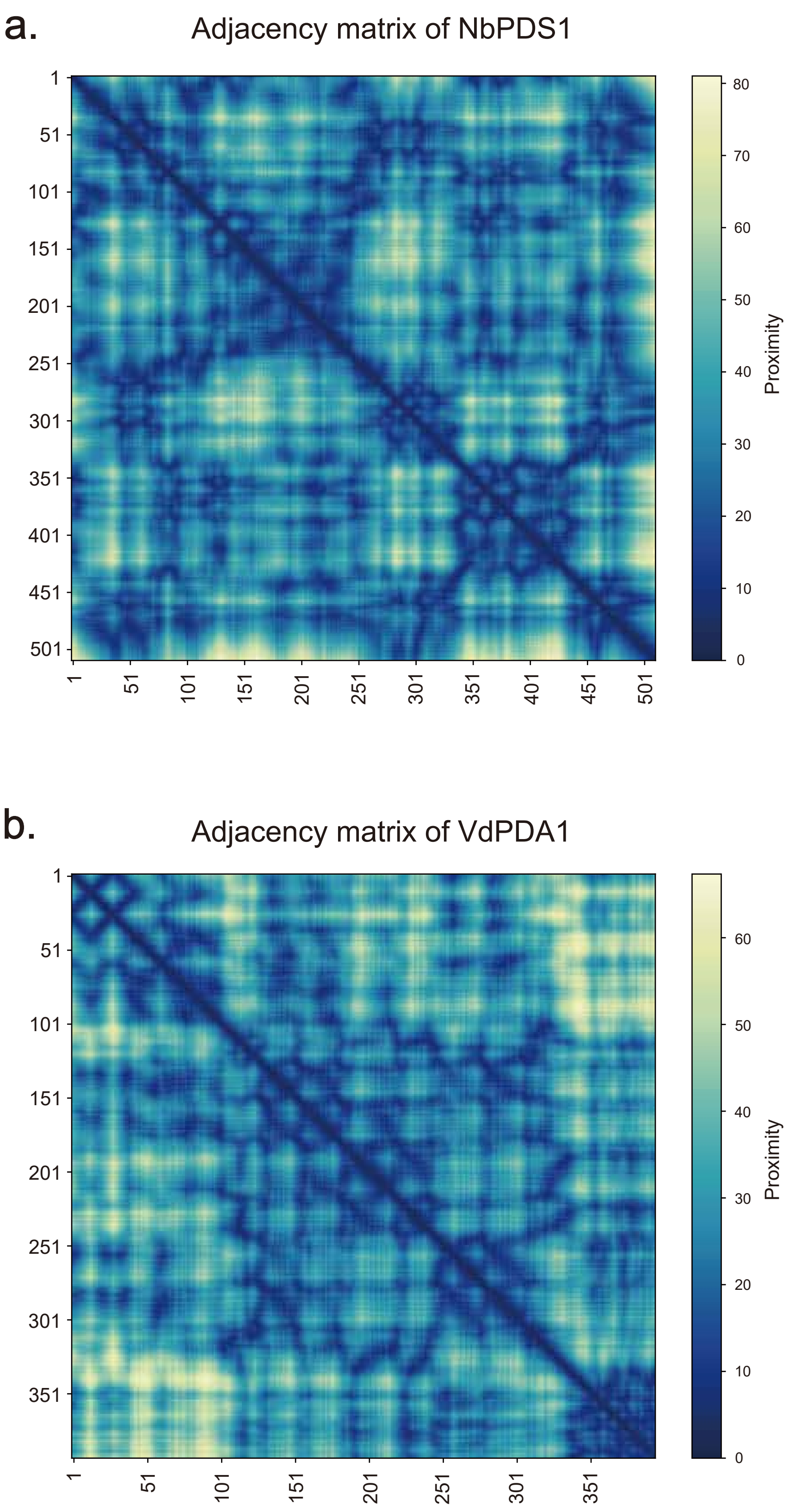

### Supplemental Figure 4

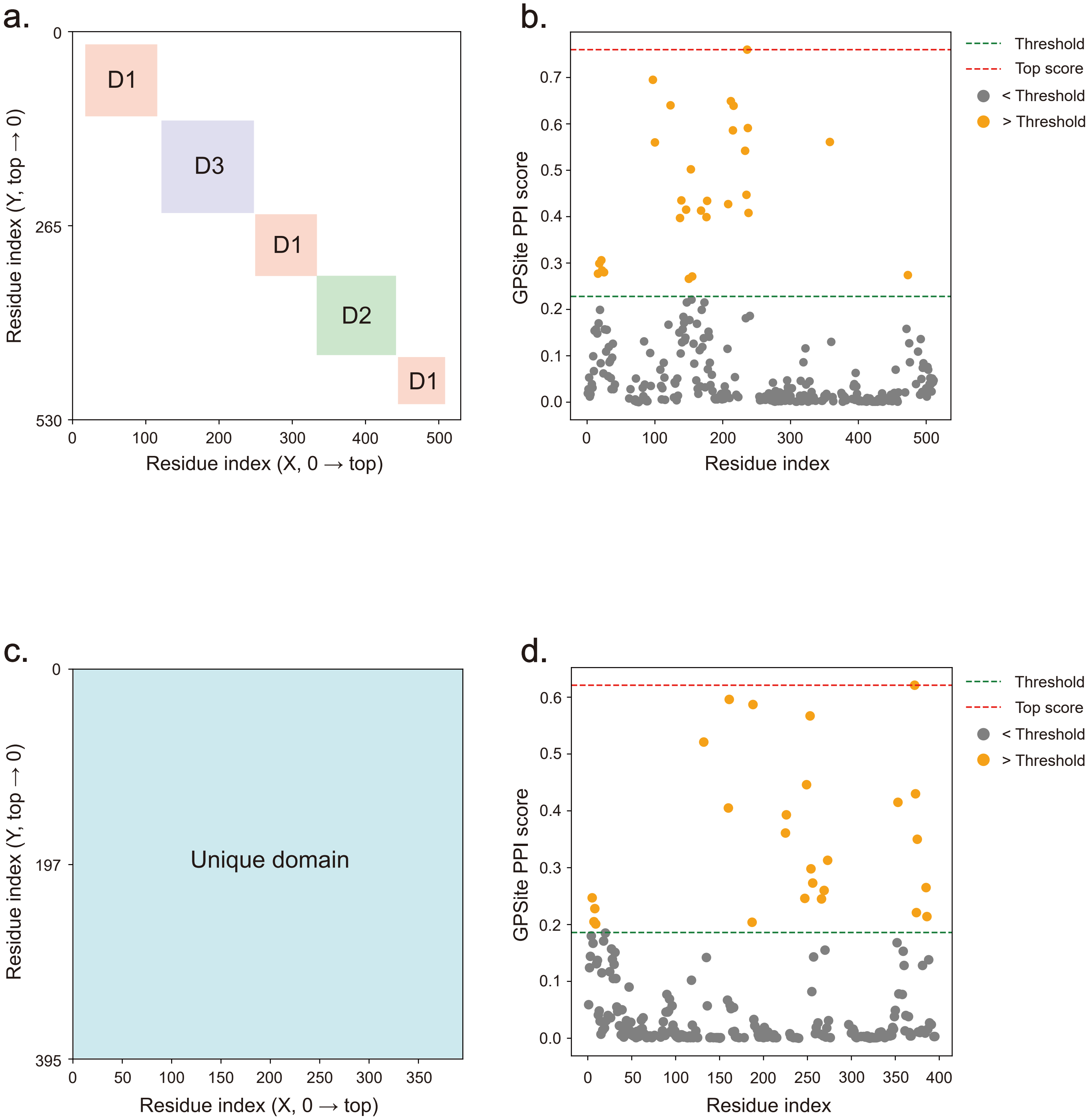
